# PhyloFold: Precise and Swift Prediction of RNA Secondary Structures to Incorporate Phylogeny among Homologs

**DOI:** 10.1101/2020.03.05.975797

**Authors:** Masaki Tagashira

**Affiliations:** Department of Computational Biology and Medical Sciences, University of Tokyo, Chiba, 277-8561, Japan; Artificial Intelligence Research Center, AIST, Tokyo, 135-0064, Japan

## Abstract

**Motivation:** The *simultaneous consideration of sequence alignment and RNA secondary structure, or structural alignment*, is known to help predict more accurate secondary structures of homologs. However, the consideration is heavy and can be done only roughly to decompose structural alignments.

**Results:** The PhyloFold method, which predicts secondary structures of homologs considering likely pairwise structural alignments, was developed in this study. The method shows the best prediction accuracy while demanding comparable running time compared to conventional methods.

**Availability:** The source code of the programs implemented in this study is available on “https://github.com/heartsh/phylofold” and “https://github.com/heartsh/phyloalifold“.

**Supplementary information:** Supplementary data are available.

## 1 Introduction

A large quantity of functional *ncRNA*s have been discovered through sequencing technology, including high throughput sequencing (Maxam and Gilbert, 1977; Bentley *et al*., 2008). The RNAs are related to various biological processes such as epigenetic silencing (Pasmant *et al*., 2011), splicing regulation (Ji *et al*., 2003), translational control (Long and Caceres, 2009), apoptosis regulation, and cell cycle control (Kino *et al*., 2010). However, the functions of a large number of RNAs have not yet been uncovered. Secondary structures of the RNAs are an important key to uncovering the functions of the RNAs because the structures play critical roles in biological processes, (Wu *et al*., 1991) and secondary structures are often conserved among homologs even in the case where the sequences of the homologs are nonconserved. (Klein and Eddy, 2003)

To more correctly predict secondary structures, simultaneous consideration of sequence alignment and secondary structure, *or structural alignment* (Sankoff, 1985; Hofacker *et al*., 2004; Havgaard *et al*., 2007), has been taken into account. (Hamada *et al*., 2009c, 2011; Tan *et al*., 2017) However, the consideration has been proven intractable (Sankoff, 1985) and thus has been conducted roughly to decompose structural alignments. (Hamada *et al*., 2009c,a; Sato *et al*., 2012; Tan *et al*., 2017) To more accurately accomplish this goal, the PhyloFold method (Figure 1) was developed in this study. The method predicts secondary structures of homologs considering likely pairwise structural alignments between the homologs by a *sparsification* technique.

**Fig. 1.**
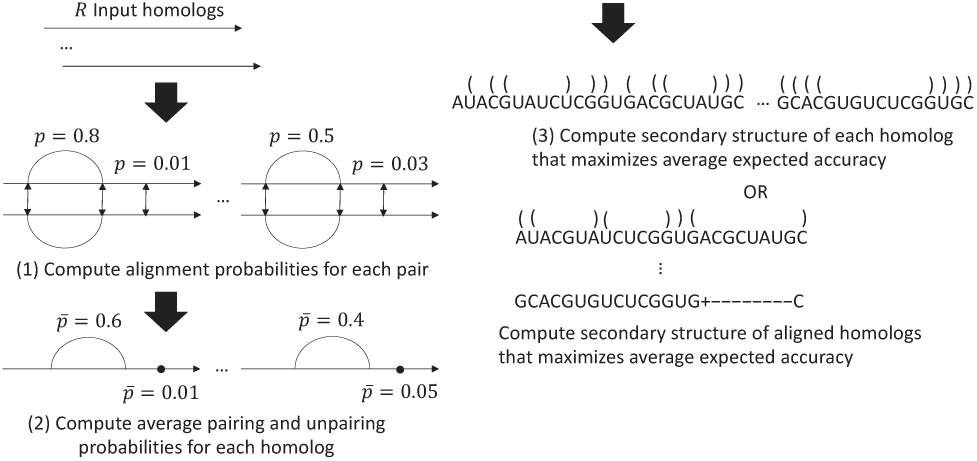
The proposed workflow of the PhyloFold method that predicts single and consensus RNA secondary structures. (1) The method computes pairwise structural alignment probabilities for each pair of input homologs. (2) Next, the method calculates average pairing probabilities between a target homolog and multiple support homologs to incorporate more than one sample into the subsequent predictions of secondary structures. (3) Finally, the method predicts the single secondary structure of each input homolog or the consensus secondary structure of the input sequence alignment.

## 2 Methods

### 2.1 Pairwise structural alignment

Let ***SS***_***RNA***_, ***SA***_ℝℕ𝔸_, 𝕊𝕋𝔸_ℝℕ𝔸_ be the (single) RNA secondary structure of the RNA sequence ***RNA***, the (pairwise) sequence alignment between the pair of RNA sequences ℝℕ𝔸, and the structural alignment between the pair ℝℕ𝔸, respectively. An alignment 𝕊𝕋𝔸_ℝℕ𝔸_ is composed of the alignment and structures ***SA***_ℝℕ𝔸_, ***SS***_***RNA***_, ***SS***_***RNA***_′, i.e. 𝕊𝕋𝔸_ℝℕ𝔸_ = (***SS***_***RNA***_, ***SS***_***RNA***_′, ***SA***_ℝℕ𝔸_) where ℝℕ𝔸 = (***RNA, RNA***′). (Figure 2) Assume secondary structures without any *pseudoknots* and *collinear* sequence alignments because their considerations lead to larger computational complexities (Sato *et al*., 2011; Bradley *et al*., 2009). Also assume structural alignments without any *indels of 2-loop*s (Supplementary section 1.1), i.e. positions must be pairing if they are aligned with another set of pairing positions. (Sankoff, 1985) Two sets of positions are said to be *pair-aligned* if they are pairing and aligned. Two positions are said to be *loop-aligned* if they are unpaired and aligned.

**Fig. 2.**
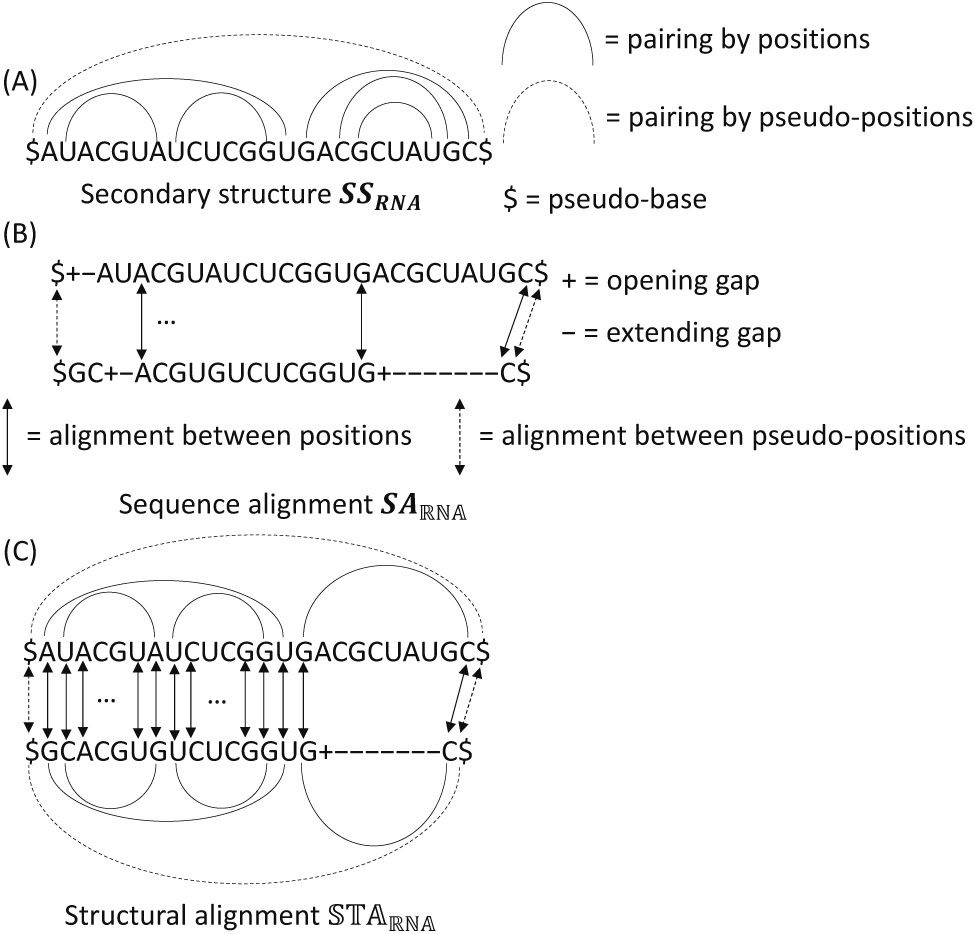
Examples of (A) a secondary structure ***SS***_***RNA***_, (B) a sequence alignment ***SA***_***RNA***_, and (C) a structural alignment 𝕊𝕋𝔸_ℝℕ𝔸_.

### 2.2 Posterior *pair alignment probability* matrix

Let 𝔖𝔗𝔄_ℝℕ𝔸_, *fe*_𝕊𝕋𝔸_ be a set of all possible alignments 𝕊𝕋𝔸_ℝℕ𝔸_ and the free energy (or inverse of the score) of the alignment 𝕊𝕋𝔸, respectively. Assume that the probability of any alignment 𝕊𝕋𝔸 ∈ 𝔖𝔗𝔄_ℝℕ𝔸_, *p*_𝕊𝕋𝔸_, obeys a *Boltzmann (probability) distribution*, i.e. 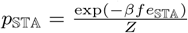 where *Z* = Σ_𝕊𝕋𝔸_ exp(− *βfe*_𝕊𝕋𝔸_) and *β* is a positive real number. *Z* is called a *partition function. β* scales any energy *fe*_𝕊𝕋𝔸_.

Let 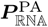 be the *pair alignment probability* matrix given the pair ℝℕ𝔸. Let 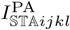 be 1 if the pairs (*i, j*), (*k, l*) are pair-aligned in the alignment 𝕊𝕋𝔸 and 0 otherwise, where *i, j* are two positions in the sequence ***RNA***, *k, l* are two positions in the other sequence ***RNA***′, and *i* < *j,k* < *l*. The matrix 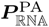 can be written by the probabilities 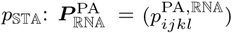 where 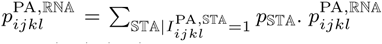 is the probability that the pairs (*i, j*), (*k, l*) are pair-aligned.

Any computation methods for a matrix 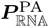 have not been proposed although the concept of it 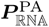 is proposed in Hamada *et al*. (2009c). Therefore, a computation method for the matrix 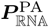 and an efficient sparse version of the method are suggested in this study.

### 2.3 Composition of structural alignment free energy *fe*_𝕊𝕋𝔸_

Let *fe*_***SS***_ be the free energy of the structure ***SS***. The energy 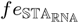 is decomposed into additional components: 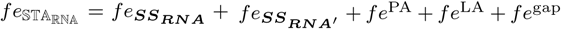. Here *fe*^PA^, *fe*^LA^ are the sum free energy of the pair and loop alignments in 𝕊𝕋𝔸_ℝℕ𝔸_, respectively. Also, *fe*^gap^ is the sum free energy (or penalty) of affine gaps in the alignment 𝕊𝕋𝔸_ℝℕ𝔸_.

The components 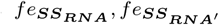 can be computed by the estimated parameters of the Turner (nearest neighbor) model, which approximates the free energy of RNA secondary structure on thermodynamics. (Turner and Mathews, 2010) As the components, conventional methods based on structural alignment employ posterior pairing probability matrices on single secondary structure, predicted by *inside-outside algorithm*s such as the McCaskill algorithm (McCaskill, 1990), to simplify computations, although the suitability of the matrices has not been discussed. (Hofacker *et al*., 2004; Do *et al*., 2008). Hence, the Turner model, instead of the matrices, is adopted in this study to prevent a reduction in the accuracy of a matrix 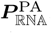.

The components *fe*^PA^, *fe*^LA^ can be computed by the RIBOSUM score matrices, predicted from learning datasets of validated structural alignments. (Klein and Eddy, 2003) In this study, the RIBOSUM score 80-65 matrices, which are the most popular, are adopted.

In the next section, an efficient (however nevertheless impractical) method that computes a matrix 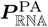 in the framework of an inside-outside algorithm is proposed.

### 2.4 Inside-outside algorithm that computes pair alignment probability matrix 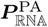

A probability 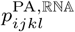 can be written with two “inside” and one “outside” partition functions 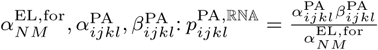 where *N, M* are the lengths of the sequences ***RNA, RNA***′, respectively. Inside and outside partition functions can be computed with those of shorter and longer substrings, respectively, and are stored in dynamic programming memory for the remaining computation. A matrix 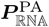 can be computed by Algorithm 1 with the time and space complexities *O*(*N* ^4^*M* ^4^), *O*(*N* ^3^*M* ^3^). Variants of Algorithm 1 with the time and space complexities *O*(*N* ^3^*M* ^3^), *O*(*N* ^2^*M* ^2^) can be considered though the sparsification proposed in this study is not effective for the variants.

Algorithm 1 is the “simultaneous” solution of the Durbin (*forward-backward*) algorithm, which predicts posterior alignment probability matrices on pairwise sequence alignment (Durbin *et al*., 1998), and the McCaskill algorithm, as expected. Algorithm 1 is also an inside-outside algorithm version of the Sankoff algorithm, which predicts pairwise structural alignments whose free energy is minimum (Sankoff, 1985), as expected. The time and space complexities of Algorithm 1 are preferably around *O*(*NM*) to deal with long ncRNAs. In the next section, a solution to the heavy complexities *O*(*N* ^4^*M* ^4^), *O*(*N* ^3^*M* ^3^), sparsifying all possible structural alignments 𝔖𝔗𝔄_ℝℕ𝔸_, is introduced.

#### Algorithm 1 An inside-outside algorithm that computes a pair alignment p robability matrix 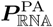.

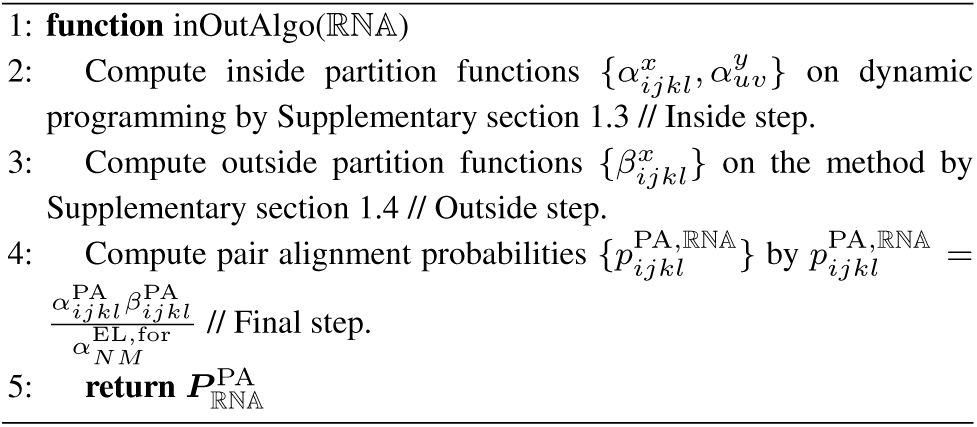

### 2.5 Sparsifying pair alignment probability matrix 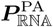

From all the possible alignments 𝔖𝔗𝔄_ℝℕ𝔸_, *sparsification* can pick out those that are favorable (e.g. with adequately low free energy). It can allow Algorithm 1 to compute only the functions 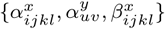 that satisfy sparsification conditions. Let 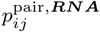 be the pairing probability of the positions *i, j* given the sequence ***RNA***. In this study, the following sparsification conditions are introduced:

- |*u* − *v*| ≤ *δ*, |(*M* − *u*) − (*N* − *v*)| ≤ *δ* for any positions *u, v*
- |(*i* − *j*) − (*k* − *l*)| ≤ *δ* for any pairs (*i, j*), (*k, l*)
- 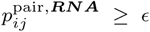 for any pairing positions *i, j* and any sequence ***RNA***

where *δ, ϵ* are sparsification parameters. The first two conditions let Algorithm 1 not consider the alignments 𝔖𝔗𝔄_ℝℕ𝔸_ with too many gaps. The last condition makes Algorithm 1 not consider the alignments 𝔖𝔗𝔄_ℝℕ𝔸_ with pairings difficult to predict (e.g. distant). Algorithm 1 with the above conditions are ideal with the time and space complexities *O*(*L*^2^), *O*(*L*^2^) where *L* = max(*N, M*) if the parameters *δ, ϵ* take sufficiently small and large values, respectively.

### 2.6 Probabilistic consistency transformation

Probabilistic consistency transformation is the technique that converts a probability between a target homolog and each support homolog into a metric that summarizes the phylogeny among all the homologs. (Do *et al*., 2005) The transformation has been required because the computational complexities involved in computing posterior probabilities among all the homologs is NP-complete as with multiple precise alignment. (Do *et al*., 2005; Hamada *et al*., 2009c; Tan *et al*., 2017) In this study, the methods that transform probabilities 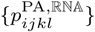 are proposed. To average probabilities 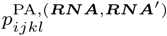 between the target homolog and each support homolog ***RNA***, {***RNA***′}, the average pairing probability 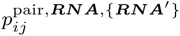 is gained:

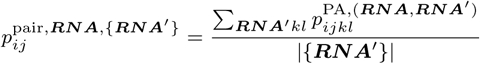

To sum probabilities 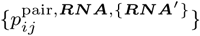, an average unpairing probability 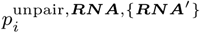 is obtained:

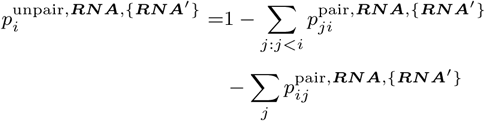

Obviously, the difference between proposed and existing probabilities 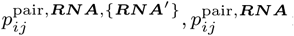 is whether multiple support homologs {***RNA***′} are considered or not.

### 2.7 Secondary structure prediction that maximizes average expected accuracy incorporating multiple support homologs

The *accuracy* of a predicted structure ***SS*** against a reference alignment 𝕊𝕋𝔸 is measured based on terms of positive and negative predictions:

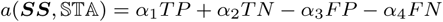

where *T P, T N, FP, FN* are the numbers of true positive, true negative, false positive, and false negative predictions, respectively, and {*α*_*h*_} are their scale parameters. In this study, the counts *T P, T N, FP, FN* are configured as:

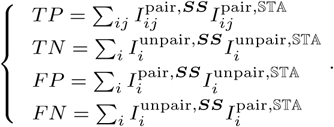

Here, 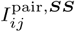 is 1 if the positions *i, j* are pairing in the structure ***SS*** and 0 otherwise; 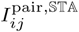 is 1 if the positions *i, j* are pairing in the alignment 𝕊𝕋𝔸_*ij*_ and 0 otherwise; 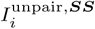 is 1 if the position *i* is unpaired in the structure ***SS*** and 0 otherwise, 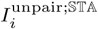 is 1 if the position *i* is unpaired in the alignment 𝕊𝕋𝔸 and 0 otherwise; and 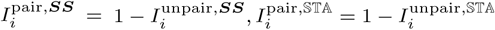

Because the accuracy and *γ*-dependent accuracy

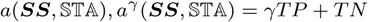

are equivalent, the expected accuracy to be maximized is gained:

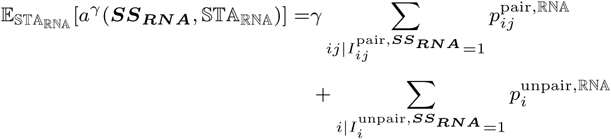

where 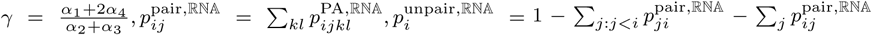. The predicted structure that maximizes expected accuracy 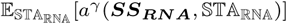 can be computed to conduct Nussinov type dynamic programming (Nussinov *et al*., 1978) based on the following recursion:

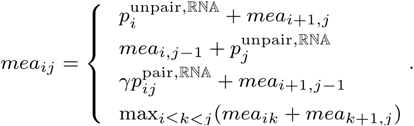

In order to consider more than one support homolog, it is sufficient that probabilities 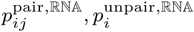 are replaced with the probabilities 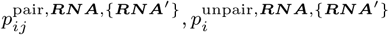 in the above recursion.

Let ***SA***_{***RNA***}_ be the sequence alignment among sequences {***RNA***}. Secondary structure prediction is extended to that of sequence alignment to view positions {*i*} on a sequence ***RNA*** as columns on an alignment ***SA***_{***RNA***}_ on the above recursion. It is known that pairing probabilities of columns *i, j* given an alignment ***SA***_{***RNA***}_, 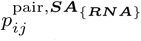, improves the prediction accuracy of consensus secondary structure. (Hamada *et al*., 2011; Bernhart *et al*., 2008) Thus, the mixture of probabilities 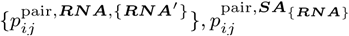, is used on the above recursion to predict the structure of an alignment ***SA***_{***RNA***}_:

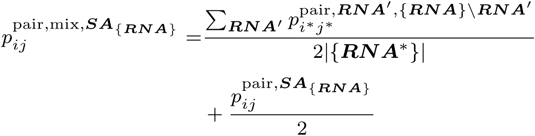

where ***RNA***′ ∈ {***RNA***}, {***RNA***\*} is the subset of the sequences {***RNA***} that is not mapped to gaps on the columns *i, j* in the alignment ***SA***_{***RNA***}_, and *i*\*, *j*\* are the positions mapped to the columns *i, j* in the alignment ***SA***_{*RNA*}_ respectively. A matrix 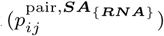 can be computed by the RNAalipfold algorithm. (Bernhart *et al*., 2008)

## Results

### 3.1 Implementations and benchmark environments

The PhyloFold method proposed in this study was implemented in the Rust programming language as the PhyloFold program (“https://github.com/heartsh/phylofold“) and the PhyloAliFold program (“https://github.com/heartsh/phyloalifold“). The programs employ multi-threading to achieve the fastest possible running time. Programs were run on a computer composed of an “AMD EPYC 7501” CPU with 64 threads and a clock rate 2GHz and 128GB of RAM unless environments for running programs are ohterwise specified.

### 3.2 Benchmark of PhyloFold method with competitors

Positive predictive value, sensitivity, and false positive rate are calculated from the numbers of true and false positives and negatives:

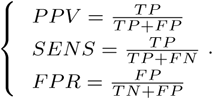

The PhyloFold program performed the best trade-off of the above metrices (Figure 3) among the PhyloFold, CentroidFold (Hamada *et al*., 2009b), CentroidHomFold (Hamada *et al*., 2009c), and TurboFold-smp (Tan *et al*., 2017) programs (Table 2) on test set “unaligned” (Table 3) while demanding comparable running time (Table 4). Ten percent of ncRNA families whose reference seed structural alignments have at most 200 columns and whose number of homologs is less than 11 are sampled from the Rfam database (Kalvari *et al*., 2018). The reference seed structural alignments of the 147 sampled families are compiled as test set “aligned”. The PhyloAliFold program demonstrated a superior trade-off of the above metrices (Figure 4) compared to the CentroidAliFold program (Hamada *et al*., 2011) on the set keeping comparable running time (Table 5).

**Table 1.**
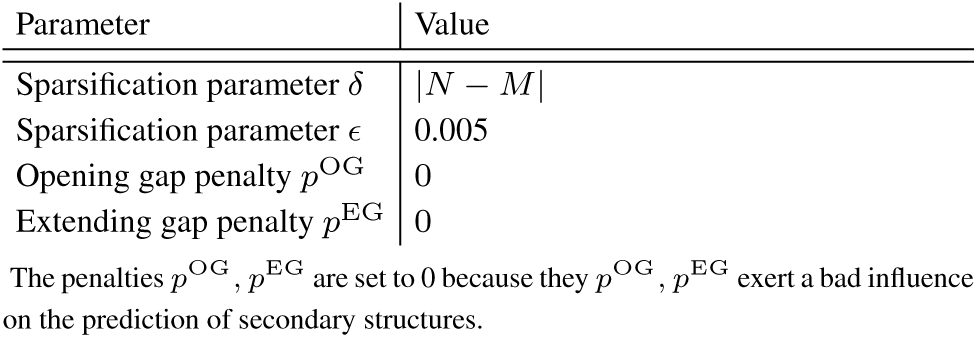
Parameter values used in the PhyloFold method.

**Table 2.**
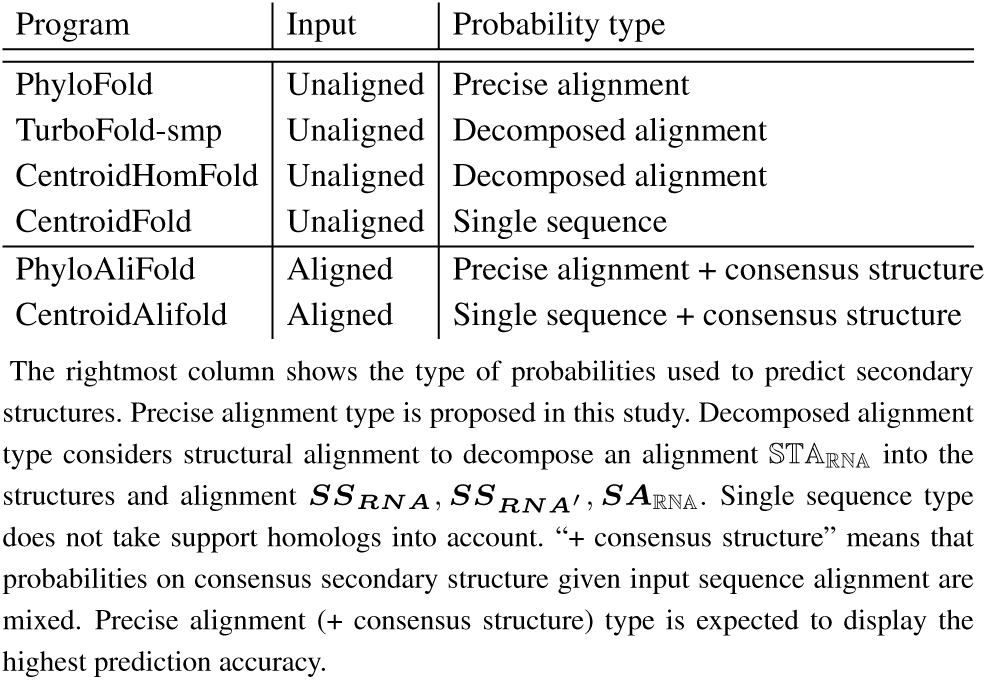
The profile of *γ*-dependent programs that predict secondary structures.

**Table 3.** The profile of test set “unaligned”.

| Family | # homologs | Range nt |
| --- | --- | --- |
| RNAIII | 4 | 489–517 |
| Hammerhead ribozyme (type III) | 10 | 40–58 |
| Hammerhead ribozyme (type I) | 10 | 45–118 |
| Bicoid 3 prime-UTR regulatory element | 4 | 543–551 |
| R2 RNA element | 10 | 202–237 |
| Gammaretrovirus core encapsidation signal | 10 | 100–102 |
| RNase MRP | 5 | 246–305 |
| Ciliate telomerase RNA | 10 | 160–215 |
| Coronavirus 3 prime stem-loop... | 10 | 43 |
| Vimentin 3 prime UTR... | 10 | 65–77 |
| RNase E 5 prime UTR element | 6 | 337–339 |

Unaligned homologs of 11 ncRNA families and the reference secondary structures of the homologs were collected from the RNA STRAND database (Andronescu *et al.*, 2008) and compiled as test set “unaligned”. The Rfam database was not used to build the set because this database does not register reference single secondary structures. Ten homologs and their reference secondary structures were sampled from each ncRNA family when it had 11 or more homologs. The middle and rightmost columns show the number of homologs and the range of lengths on each ncRNA family, respectively. “Coronavirus 3 prime stem-loop...” and “Vimentin 3 prime UTR” stand for “Coronavirus 3 prime stem-loop II-like motif (s2m)” and “Vimentin 3 prime UTR protein-binding region,” respectively.

**Table 4.** The benchmark of *γ*-dependent programs that predict single secondary structures on running time using test set “unaligned”.

| Program (algorithm) | Running time | Time complexity |
| --- | --- | --- |
| PhyloFold | 4.95 m | $O(R^2 N^2 + RN^3)$ |
| TurboFold-smp | 4.48 m | $O(R^2 N^2 + RN^3)$ |
| CentroidHomFold | 1.61 m | $O(R^2 N^3)$ |
| CentroidFold (CentroidFold) | 8.26 s | $O(RN^3)$ |
| CentroidFold (CONTRAfold) | 8.31 s | $O(RN^3)$ |

**Table 5.** The benchmark of *γ*-dependent programs that predict consensus secondary structures on running time using test set “aligned”.

| Program | Running time | Time complexity |
| --- | --- | --- |
| PhyloAliFold | 58.3 s | $O(R^2 N^2 + RC^2 + C^3)$ |
| CentroidAliFold | 1.06 s | $O(RN^3 + RC^2 + C^3)$ |

**Fig. 3.**
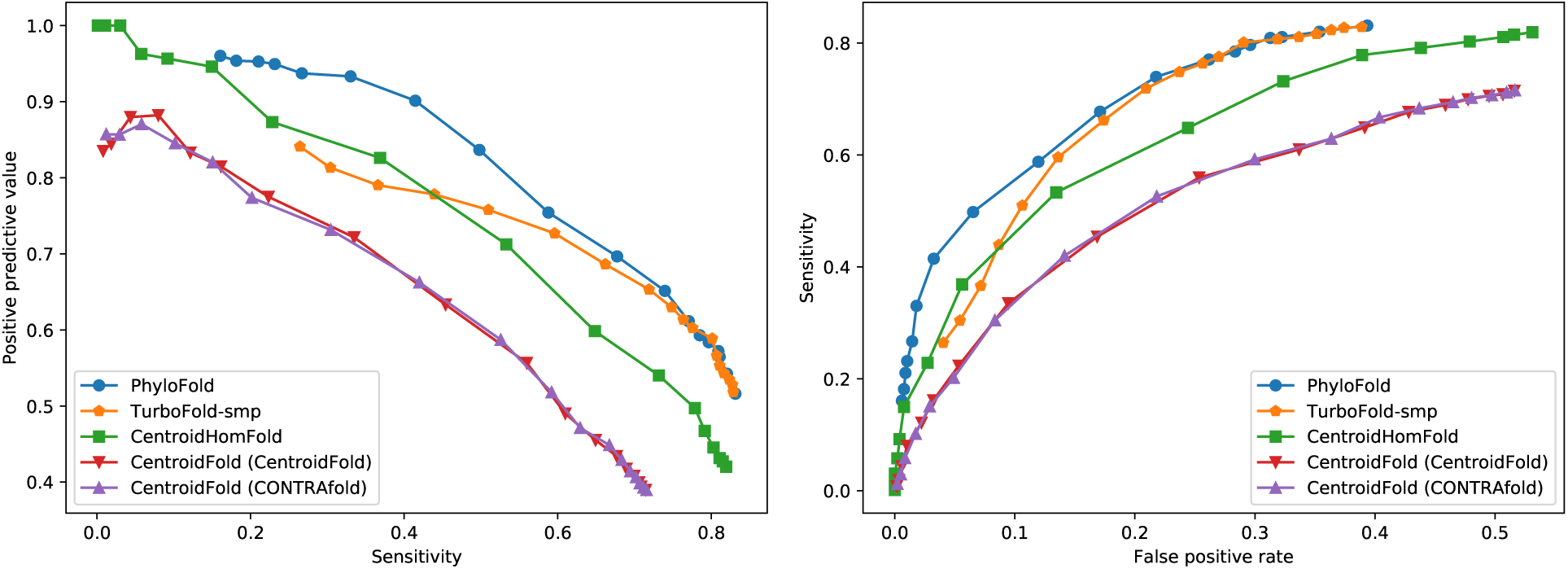
The benchmark of *γ*-dependent programs that predict single secondary structures on prediction accuracy using test set “unaligned”. The trade-off curves are composed of (left) pairs {(*SENS, PPV*)} and (right) pairs {(*FPR, SENS*)} at each parameter *γ* = 2^*g*^ : *g* ∈ {−7, …, 10} respectively. The PhyloFold method specified its parameter values as in Table 1. The CentroidFold and CentroidHomFold programs were run with their default parameter values. The TurboFold-smp was given only the number of its iterations *η* = 1. The different algorithms are shown in parentheses.

**Fig. 4.**
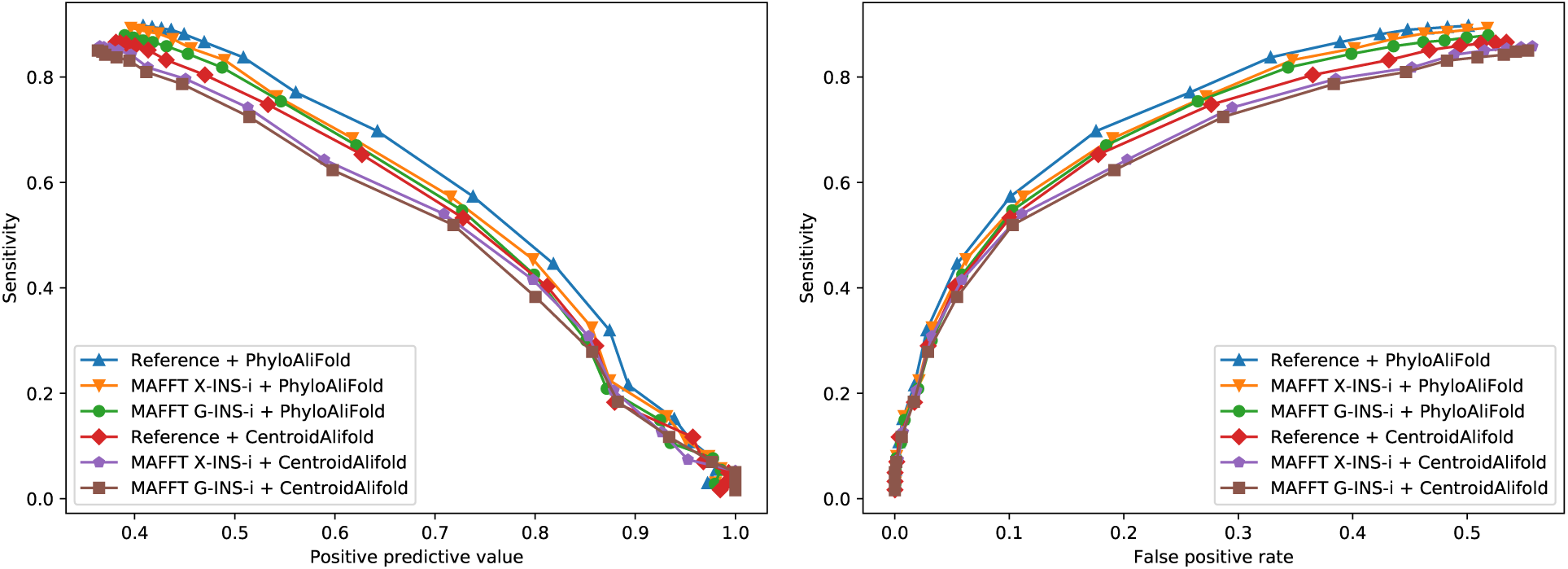
The benchmark of *γ*-dependent programs that predict consensus secondary structures on prediction accuracy using test set “aligned”. The trade-off curves are composed of (left) pairs {(*PPV, SENS*)} and (right) pairs {(*FPR, SENS*)} at each parameter *γ* = 2^*g*^ respectively. The parameter values of the PhyloFold method are the same as Figure 3. The CentroidAlifold program was run with its default parameter values. The MAFFT G-INS-i and MAFFT X-INS-i programs (Katoh and Standley, 2013) were used to compute the sequence alignments to input into the PhyloAliFold program. The MAFFT G-INS-i executes only classic multiple sequence alignment whereas the MAFFT X-INS-i considers structural alignments during multiple sequence alignment to decompose them. The reference sequence alignments of the ncRNA families in test set “aligned” are also used as the input of the PhyloAliFold program.

## 4 Conclusions

The PhyloFold method, which predicts secondary structures considering the phylogeny among homologs, has been devised. The method demonstrated superior prediction accuracy compared to competitors while demanding comparable running time on the prediction of both single and consensus secondary structures. The method has the potential to be improved by secondary structure probing methods such as icSHAPE (Spitale *et al*., 2015) and SHAPE-MaP (Siegfried *et al*., 2014) as demonstrated in the SuperFold (Siegfried *et al*., 2014) and TurboFold methods.

Probabilities 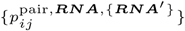 can be used to measure the reliability of predicted secondary structures as a part of postprocessing. (Figure 5) Also, the probabilities 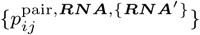can be utilized to improve the accuracy of existing methods that predict RNA-RNA interactions, alignments, etc., such as the IntaRNA (Mann *et al*., 2017), CentroidAlign (Hamada *et al*., 2009a), and DAFS (Sato *et al*., 2012) programs, which either employ the decomposition of structural alignments or do not incorporate structural alignments.

**Fig. 5.**
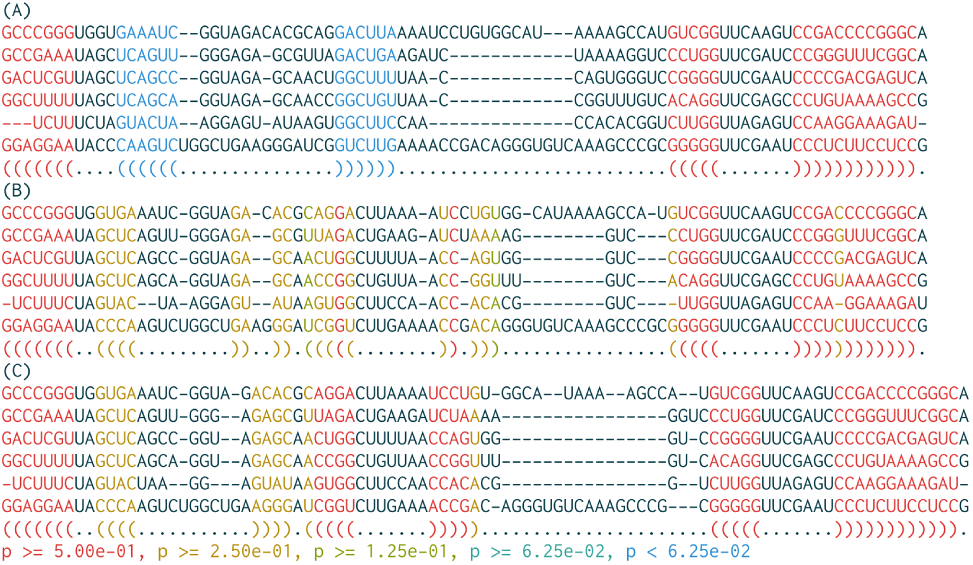
Consensus secondary structures color-coded with their reliability. The structures are of the sequence alignment among homologs sampled from the tRNA family registered in the Rfam database. The alignment was computed by (A) the MAFFT G-INS-i and (B) MAFFT X-INS-i programs. The structures were computed by the RNAalifold program (Bernhart et al., 2008). (C) The reference consensus secondary structures of the homologs derived from the database. The reliability was assessed to average mean pairing probabilities 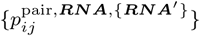 per column pair. The reliability was improved when the quality of the input sequence alignment increased.

## Acknowledgements

I thank the members of the Asai and Frith laboratories for discussing this study with me for years. Also, I thank Dr. Risa Kawaguchi for sharing valuable information about the scaling and logsumexp methods on exact probability computations with me. I would like to thank Editage (www.editage.com) for English language editing. Most computations were performed on the NIG supercomputer at the ROIS National Institute of Genetics.

## Author contributions

M.T. conceived the proposed method, implemented it in the presented programs, experimented with them and competitors, and wrote this paper.

