## Supplementary materials for "PhyloFold: Precise and Swift Prediction of RNA Secondary Structures to Incorporate Phylogeny among Homologs"

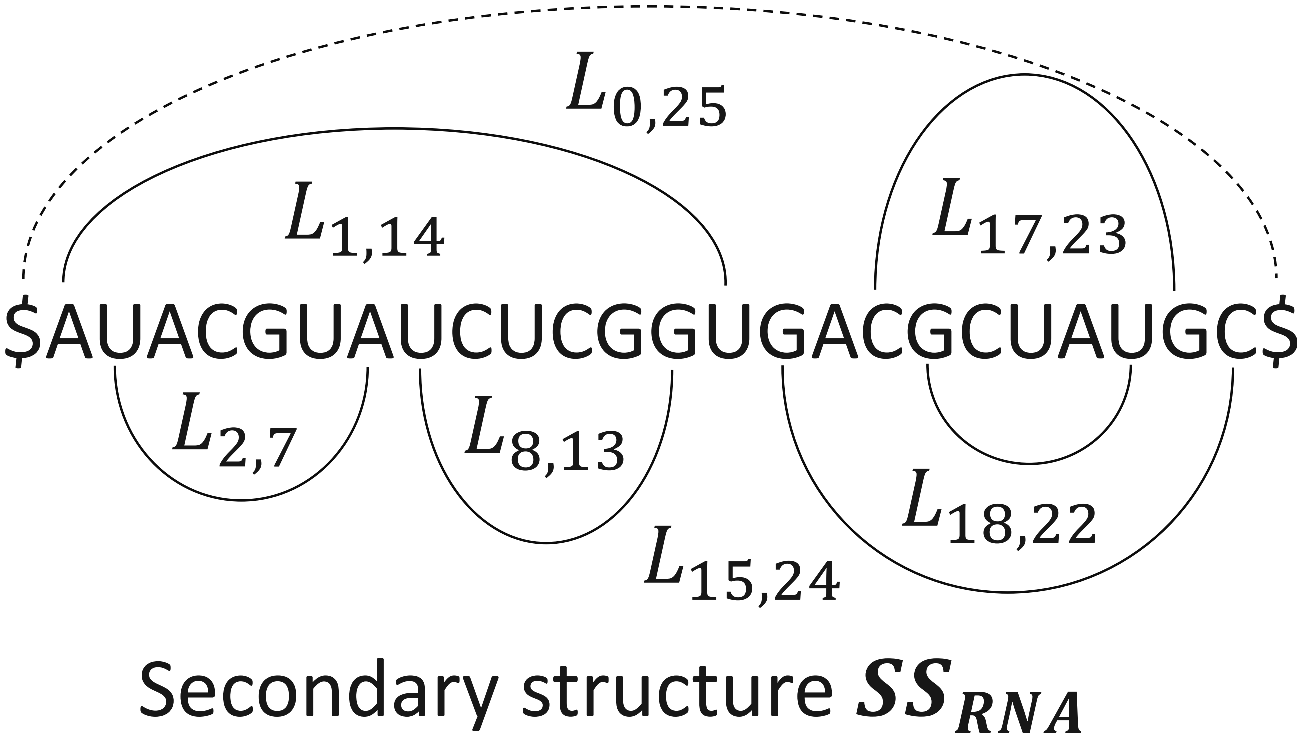


**Fig. 1.** A secondary structure $\boldsymbol{SS_{RNA}}$ with nested loops. $L_{0,25} = \{1, 14, 15, 24\}, L_{1,14} = \{2, 7, 8, 13\}$ are external and internal 3-loops, respectively. $L_{15,24} = \{16, 17, 23\}, L_{17,23} = \{18, 22\}$ are internal 2-loops. $L_{2,7} = \{3, \ldots, 6\}, L_{8,13} = \{9, \ldots, 12\}, L_{18,22} = \{19 \ldots, 21\}$ are internal 1-loops.

### 1 Recursions of partition functions $\{\alpha^x_{ijkl}, \alpha^y_{uv}, \beta^x_{ijkl}\}$ for Algorithm 1

#### 1.1 RNA secondary structures with loops on Turner model

A position $u$ is said to be *accessible from pairing* (or *closed by*) positions $i, j$ if $i < u < j$ and the position pair $(i, j)$ is the closest to $u$ of all pairing position pairs. A position set about pairing positions $i, j$, $L_{ij} = \{u | I^{\mathrm{loop}}_{uij} = 1\}$, is the *loop* of them $i, j$ where $I^{\mathrm{l}}_{uij}$ is 1 if the position $u$ is accessible from them $i, j$ and 0 otherwise. A loop $L_{ij}$ is a *b-loop* if the loop $L_{ij}$ contains $b - 1$ pairing position pairs accessible from the positions $i, j$. $b$-loop is divided into three classes on the Turner model: 1-loop (= *hairpin loop*), 2-loop (= *stem, bulge, and interior loop*), and $(b > 2)$-loop (= *multi-loop*). A loop $L_{ij}$ is said to be *internal* or *external* if the positions $i, j$ are accessible from loops of other positions and pseudo-positions, respectively. RNA secondary structures can be ornamented with nested loops. (Supplementary figure 1)

On the Turner model, energy $fe_{\boldsymbol{SS}}$ is decomposed into four categories of additional component: $fe_{\boldsymbol{SS}} = \sum_{ij\lambda | I^{\mathrm{pair}, \boldsymbol{SS}}_{ij} I^{\lambda}_{ij} = 1} fe^{\lambda}_{ij}$ where $\lambda \in \{\mathrm{I1L}, \mathrm{I2L}, \mathrm{IML}, \mathrm{EL}\}$, $\{I^{\lambda}_{ij}\}$ are 1 if the loop $L_{ij}$ is an internal 1-loop, an internal 2-loop, an internal multi-loop, or an external loop, respectively, and $\{fe^{\lambda}_{ij}\}$ is the free energy of the loop $L_{ij}$ when the loop $L_{ij}$ is the four classes of loop, respectively. Energy $fe^{\mathrm{I2L}}_{ij}$ depends on the pairing positions $m, n$ accessible from the positions $i, j$: $fe^{\mathrm{I2L}}_{ij} = fe^{\mathrm{I2L}}_{ijmn}$ where $fe^{\mathrm{I2L}}_{ijmn}$ is the energy $fe^{\mathrm{I2L}}_{ij}$ parameterized with the positions $m, n$. The Turner model restricts the number of unpaired positions of the 2-loop $L_{ij}$, $(m-i)+(j-n)+2$: $(m-i)+(j-n)+2 \leq 30$ to reduce time complexities of prediction algorithms. Energy $fe^{\mathrm{IML}}_{ij}$ is decomposed into the terms of accessible and closing pairings: $fe^{\mathrm{IML}}_{ij} = fe^{\mathrm{IML,CBP}} + \sum_{mn | m,n \in L_{ij}} fe^{\mathrm{IML,ABP}}$ where $fe^{\mathrm{IML,CBP}}$, $fe^{\mathrm{IML,ABP}}$ are free energy per closing and accessible pairing. Energy $fe^{\mathrm{EL}}_{ij}$ does not influence the entire free energy: $fe^{\mathrm{EL}}_{ij} = 0$. All the above energy $\{fe^{\mathrm{I1L}}_{ij}, fe^{\mathrm{I2L}}_{ijmn}, fe^{\mathrm{IML,CBP}}, fe^{\mathrm{IML,ABP}}\}$ is available as the different parameter sets of the Turner model (Turner and Mathews, 2010). The Turner 2004 model, the updated version of the Turner 1999 model, was chosen in this study.

#### 1.2 Detail of alignment score

The components $fe^{\mathrm{PA}}, fe^{\mathrm{LA}}$ can be computed to sum RIBOSUM scores across all pair-aligned and loop-aligned positions respectively:

$$\begin{cases} fe^{\mathrm{PA}} = \sum_{ijkl | I^{\mathrm{PA}, \mathbb{STA}}_{ijkl}} \ln \frac{p^{\mathrm{PA}}_{ijkl}}{p^{\mathrm{PA,rand}}_{ijkl}} \\ fe^{\mathrm{LA}} = \sum_{uv | I^{\mathrm{uv}, \mathbb{STA}}_{uv}} \ln \frac{p^{\mathrm{LA}}_{uv}}{p^{\mathrm{LA,rand}}_{uv}} \end{cases}$$

where $p^{\mathrm{PA}}_{ijkl}, p^{\mathrm{PA,rand}}_{ijkl}$ are the *prior* pair alignment probabilities of the pairs $(i, j), (k, l)$ in the assumptions whose pair $\mathbb{RNA}$ is correlated or random, respectively, $I^{\mathrm{LA}, \mathbb{STA}}_{uv}$ is 1 if the positions $u, v$ are loop-aligned in the alignment $\mathbb{STA}$ and 0 otherwise, and $p^{\mathrm{LA}}_{uv}, p^{\mathrm{LA,rand}}_{uv}$ are the prior loop alignment probabilities of the positions $u, v$ in the assumptions whose pair $\mathbb{RNA}$ is correlated or random, respectively. The penalty of a gap whose length is $G$ can be computed to sum one opening gap penalty and $G - 1$ extending gap penalties: $p^{\mathrm{OG}} + (G-1)p^{\mathrm{EG}}$ where $p^{\mathrm{OG}}, p^{\mathrm{EG}}$ are the penalties of opening and

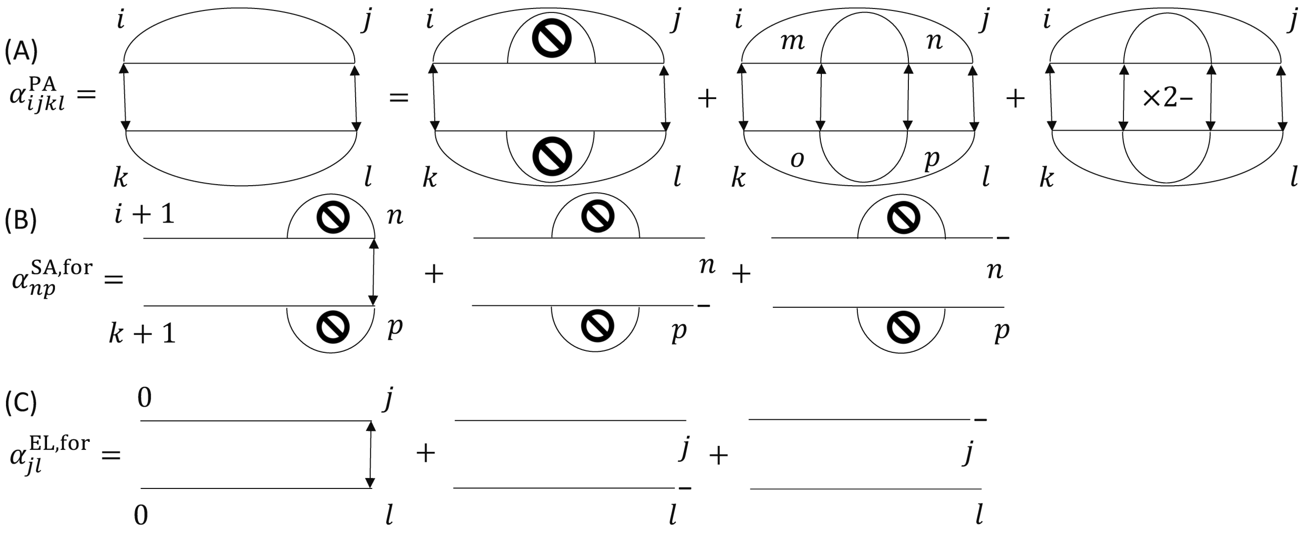


**Fig. 2.** Interpretations of the formulations of inside partition functions (A)$\alpha^{\mathrm{PA}}_{ijkl}$, (B)$\alpha^{\mathrm{SA}}_{np}$, (C)$\alpha^{\mathrm{EL}}_{jl}$.

extending gaps, respectively. In the next section, recursions to draw efficient algorithms that compute inside partition functions $\{\alpha^{x}_{ijkl}, \alpha^{y,(i,j),(k,l)}_{np}\}$ are discussed.

### 1.3 Recursions for efficient computations of inside partition functions $\{\alpha^{x}_{ijkl}, \alpha^{y}_{uv}\}$

Let $\alpha^{\mathrm{PA}}_{ijkl}$ be the inside partition function between the pair-aligned pairs $(i,j),(k,l)$. It $\alpha^{\mathrm{PA}}_{ijkl}$ consists of three additional components: $\alpha^{\mathrm{PA}}_{ijkl} = \alpha^{\mathrm{PA,I1L}}_{ijkl} + \alpha^{\mathrm{PA,I2L}}_{ijkl} + \alpha^{\mathrm{PA,IML}}_{ijkl}$ where $\alpha^{\mathrm{PA,I1L}}_{ijkl}, \alpha^{\mathrm{PA,I2L}}_{ijkl}, \alpha^{\mathrm{PA,IML}}_{ijkl}$ are the function $\alpha^{\mathrm{PA}}_{ijkl}$ whose loops $L_{ij}, L_{kl}$ are internal 1-loops, internal 2-loops, and internal multi-loops, respectively. ((A) in Supplementary figure 2) The first term $\alpha^{\mathrm{PA,I1L}}_{ijkl}$ recurses into its inner partition function: $\alpha^{\mathrm{PA,I1L}}_{ijkl} = \exp(\beta(\frac{p^{\mathrm{PA}}_{ijkl}}{p^{\mathrm{PA,rand}}_{ijkl}} - fe^{\mathrm{I1L}}_{ij} - fe^{\mathrm{I1L}}_{kl}))\alpha^{\mathrm{SA,for},(i,j),(k,l)}_{j-1,l-1}$ and $\alpha^{\mathrm{SA,for},(i,j),(k,l)}_{np} = \alpha^{\mathrm{SA,for}}_{np}$ is the partition function between the pairs $(i+1,n),(k+1,p)$ whose loops $L_{ij}, L_{kl}$ are internal 1-loops. Similarly, the remaining terms $\alpha^{\mathrm{PA,I2L}}_{ijkl}, \alpha^{\mathrm{PA,IML}}_{ijkl}$ recurse into their inner partition functions:

$$\begin{cases} \alpha^{\mathrm{PA,I2L}}_{ijkl} = \exp(\beta\frac{p^{\mathrm{PA}}_{ijkl}}{p^{\mathrm{PA,rand}}_{ijkl}})\sum_{mnop:i<m<n<j,k<o<p<l}\exp(-\beta(fe^{\mathrm{I2L}}_{ijmn} + fe^{\mathrm{I2L}}_{klop}))\alpha^{\mathrm{SA,for},(i,j),(k,l)}_{m-1,o-1}\alpha^{\mathrm{PA}}_{mnop}\alpha^{\mathrm{SA,back},(i,j),(k,l)}_{n+1,p+1} \\ \alpha^{\mathrm{PA,IML}}_{ijkl} = \exp(\beta(\frac{p^{\mathrm{PA}}_{ijkl}}{p^{\mathrm{PA,rand}}_{ijkl}} - 2fe^{\mathrm{IML,CBP}}))\alpha^{\mathrm{IML,for},(i,j),(k,l)}_{j-1,l-1} \end{cases}$$

where $\alpha^{\mathrm{SA,back},(i,j),(k,l)}_{mo} = \alpha^{\mathrm{SA,back}}_{mo}$ is the partition function between the pairs $(m,j-1),(o,l-1)$ whose loops $L_{ij}, L_{kl}$ are internal 1-loops and $\alpha^{\mathrm{IML,for},(i,j),(k,l)}_{np} = \alpha^{\mathrm{IML,for}}_{np}$ is the partition function between the pairs $(i+1,n),(k+1,p)$ whose loops $L_{ij}, L_{kl}$ are internal multi-loops.

An inner partition function $\alpha^{\mathrm{SA,for}}_{np}$ is composed of three terms: $\alpha^{\mathrm{SA,for}}_{np} = \alpha^{\mathrm{SA,for,LA}}_{np} + \alpha^{\mathrm{SA,for,gap}}_{np} + \alpha^{\mathrm{SA,for,gap\prime}}_{np}$ where $\alpha^{\mathrm{SA,for,LA}}_{np}, \alpha^{\mathrm{SA,for,gap}}_{np}, \alpha^{\mathrm{SA,for,gap\prime}}_{np}$ are the function $\alpha^{\mathrm{SA,for}}_{np}$ in the cases in which the positions $n,p$ are aligned, the position $n$ is aligned with a gap behind the position $p$, and the position $p$ is aligned with a gap behind the position $n$, respectively. ((B) in Supplementary figure 2) The functions $\alpha^{\mathrm{SA,for,LA}}_{np}, \alpha^{\mathrm{SA,for,gap}}_{np}, \alpha^{\mathrm{SA,for,gap\prime}}_{np}$ recurse into their forward partition functions:

$$\begin{cases} \alpha^{\mathrm{SA,for,LA}}_{np} = \alpha^{\mathrm{SA,for}}_{n-1,p-1}\exp(\beta\frac{p^{\mathrm{LA}}_{np}}{p^{\mathrm{LA,rand}}_{np}}) \\ \alpha^{\mathrm{SA,for,gap}}_{np} = (\alpha^{\mathrm{SA,for,LA}}_{n-1,p} + \alpha^{\mathrm{SA,for,gap\prime}}_{n-1,p})\exp(\beta p^{\mathrm{OG}}) + \alpha^{\mathrm{SA,for,gap}}_{n-1,p}\exp(\beta p^{\mathrm{EG}}) \\ \alpha^{\mathrm{SA,for,gap\prime}}_{np} = (\alpha^{\mathrm{SA,for,LA}}_{n,p-1} + \alpha^{\mathrm{SA,for,gap}}_{n,p-1})\exp(\beta p^{\mathrm{OG}}) + \alpha^{\mathrm{SA,for,gap\prime}}_{n,p-1}\exp(\beta p^{\mathrm{EG}}) \end{cases}.$$

Similarly, an inner partition function $\alpha^{\mathrm{IML,for}}_{np}$ consists of three terms: $\alpha^{\mathrm{IML,for}}_{np} = \alpha^{\mathrm{IML,for,BA}}_{np} + \alpha^{\mathrm{IML,for,gap}}_{np} + \alpha^{\mathrm{IML,for,gap\prime}}_{np}$ where $\alpha^{\mathrm{IML,for,BA}}_{np}, \alpha^{\mathrm{IML,for,gap}}_{np}, \alpha^{\mathrm{IML,for,gap\prime}}_{np}$ are the function $\alpha^{\mathrm{IML,for}}_{np}$ in the cases in which the positions $n,p$ are aligned, the position $n$ is aligned with a gap behind the position $p$, and the position $p$ is aligned with a gap behind the position $n$, respectively. The functions $\alpha^{\mathrm{IML,for,BA}}_{np}, \alpha^{\mathrm{IML,for,gap}}_{np}, \alpha^{\mathrm{IML,for,gap\prime}}_{np}$ recurse into their forward partition functions:

$$\begin{cases} \alpha^{\mathrm{IML,for,BA}}_{np} = \sum_{mo:m<n,o<p}(\alpha^{\mathrm{1P,for}}_{m-1,o-1} + \alpha^{\mathrm{IML,for}}_{m-1,o-1})\exp(-2\beta fe^{\mathrm{IML,ABP}})\alpha^{\mathrm{PA}}_{mnop} + \alpha^{\mathrm{IML,for}}_{n-1,p-1}\exp(\beta\frac{p^{\mathrm{LA}}_{np}}{p^{\mathrm{LA,rand}}_{np}}) \\ \alpha^{\mathrm{IML,for,gap}}_{np} = (\alpha^{\mathrm{IML,for,BA}}_{n-1,p} + \alpha^{\mathrm{IML,for,gap\prime}}_{n-1,p})\exp(\beta p^{\mathrm{OG}}) + \alpha^{\mathrm{IML,for,gap}}_{n-1,p}\exp(\beta p^{\mathrm{EG}}) \\ \alpha^{\mathrm{IML,for,gap\prime}}_{np} = (\alpha^{\mathrm{IML,for,BA}}_{n,p-1} + \alpha^{\mathrm{IML,for,gap}}_{n,p-1})\exp(\beta p^{\mathrm{OG}}) + \alpha^{\mathrm{IML,for,gap\prime}}_{n,p-1}\exp(\beta p^{\mathrm{EG}}) \end{cases}$$

where $\alpha^{\mathrm{1P,for}}_{np}$ is the function $\alpha^{\mathrm{IML,for}}_{np}$ whose loops $L_{ij}, L_{kl}$ contain one pairing between the pairs $(i+1,n),(k+1,p)$, respectively. The function $\alpha^{\mathrm{1P,for}}_{np}$ can be computed in a similar manner to the function $\alpha^{\mathrm{IML,for}}_{np}$.

Let $\alpha^{\mathrm{EL,for},(0,N+1),(0,M+1)}_{jl} = \alpha^{\mathrm{EL}}_{jl}$ be the inside partition function between the pairs $(0,j),(0,l)$ whose loops $L_{0,N+1}, L_{0,M+1}$ are external loops. The function $\alpha^{\mathrm{EL,for}}_{jl}$ is composed of three terms: $\alpha^{\mathrm{EL,for}}_{jl} = \alpha^{\mathrm{EL,for,BA}}_{jl} + \alpha^{\mathrm{EL,for,gap}}_{jl} + \alpha^{\mathrm{EL,for,gap\prime}}_{jl}$ where $\alpha^{\mathrm{EL,for,BA}}_{jl}, \alpha^{\mathrm{EL,for,gap}}_{jl}, \alpha^{\mathrm{EL,for,gap\prime}}_{jl}$ are the function $\alpha^{\mathrm{EL,for}}_{jl}$ in the cases in which the positions $j,l$ are aligned, the position $j$ is aligned with a gap behind the position $l$, and the position $l$ is aligned with a gap behind the position $j$, respectively. ((C) in Supplementary figure 2) The functions $\alpha^{\mathrm{EL,for,BA}}_{np}, \alpha^{\mathrm{EL,for,gap}}_{np}, \alpha^{\mathrm{EL,for,gap\prime}}_{np}$ recurse into their forward partition functions:

$$\begin{cases} \alpha^{\mathrm{EL,for,BA}}_{jl} = \sum_{ik} \alpha^{\mathrm{EL,for}}_{i-1,k-1} \alpha^{\mathrm{PA}}_{ijkl} + \alpha^{\mathrm{EL,for}}_{j-1,l-1} \exp(\beta \frac{p^{\mathrm{LA}}_{jl}}{p^{\mathrm{LA,rand}}_{jl}}) \\ \alpha^{\mathrm{EL,for,gap}}_{jl} = (\alpha^{\mathrm{EL,for,BA}}_{j-1,l} + \alpha^{\mathrm{EL,for,gap\prime}}_{j-1,l}) \exp(\beta p^{\mathrm{OG}}) + \alpha^{\mathrm{EL,for,gap}}_{j-1,l} \exp(\beta p^{\mathrm{EG}}) \\ \alpha^{\mathrm{EL,for,gap\prime}}_{jl} = (\alpha^{\mathrm{EL,for,BA}}_{j,l-1} + \alpha^{\mathrm{EL,for,gap}}_{j,l-1}) \exp(\beta p^{\mathrm{OG}}) + \alpha^{\mathrm{EL,for,gap\prime}}_{j,l-1} \exp(\beta p^{\mathrm{EG}}) \end{cases}$$

"Backward" inside partition functions $\{\alpha^{x,\mathrm{back},y}_{ijkl}, \alpha^{z,\mathrm{back},w}_{uv}\}$ are computed in a similar way to "forward" inside partition functions $\{\alpha^{x,\mathrm{for},y}_{ijkl}, \alpha^{z,\mathrm{for},w}_{uv}\}$, for example:

$$\begin{cases} \alpha^{\mathrm{SA,back},(i,j),(k,l)}_{mo} = \alpha^{\mathrm{SA,back}}_{mo} = \alpha^{\mathrm{SA,back,LA}}_{mo} + \alpha^{\mathrm{SA,back,gap}}_{mo} + \alpha^{\mathrm{SA,back,gap\prime}}_{mo} \\ \alpha^{\mathrm{SA,back,LA}}_{mo} = \exp(\beta \frac{p^{\mathrm{LA}}_{mo}}{p^{\mathrm{LA,rand}}_{mo}}) \alpha^{\mathrm{SA,back}}_{m+1,o+1} \\ \alpha^{\mathrm{SA,back,gap}}_{mo} = \exp(\beta p^{\mathrm{OG}})(\alpha^{\mathrm{SA,back,LA}}_{m+1,p} + \alpha^{\mathrm{SA,back,gap\prime}}_{m+1,p}) + \exp(\beta p^{\mathrm{EG}}) \alpha^{\mathrm{SA,back,gap}}_{m+1,o} \\ \alpha^{\mathrm{SA,back,gap\prime}}_{mo} = \exp(\beta p^{\mathrm{OG}})(\alpha^{\mathrm{SA,back,LA}}_{m,o+1} + \alpha^{\mathrm{SA,back,gap}}_{m,o+1}) + \exp(\beta p^{\mathrm{EG}}) \alpha^{\mathrm{SA,back,gap\prime}}_{m,o+1} \end{cases}.$$

To begin dynamic programming, the initial condition

$$\alpha^{\mathrm{SA,for}}_{ik} = \alpha^{\mathrm{SA,for,LA}}_{ik} = \alpha^{\mathrm{SA,back}}_{jl} = \alpha^{\mathrm{SA,back,LA}}_{jl} = \alpha^{\mathrm{EL,for}}_{0,0} = \alpha^{\mathrm{EL,for,BA}}_{0,0} = \alpha^{\mathrm{EL,back}}_{N+1,M+1} = \alpha^{\mathrm{EL,back,BA}}_{N+1,M+1} = 1$$

is used. (The other inside partition functions $\{\alpha^{x}_{ijkl}, \alpha^{y}_{uv}\}$ are set to 0.) $Z$ is gained from the equation $Z = \alpha^{\mathrm{EL,for}}_{NM}$.

#### 1.4 Recursions for efficient computations of outside partition functions $\{\beta^{x}_{ijkl}\}$

Let $\beta^{\mathrm{PA}}_{ijkl}$ be the outside partition function between the pair-aligned pairs $(i,j),(k,l)$. It $\beta^{\mathrm{PA}}_{ijkl}$ consists of three additional components: $\beta^{\mathrm{PA}}_{ijkl} = \beta^{\mathrm{PA,EL}}_{ijkl} + \beta^{\mathrm{PA,I2L}}_{ijkl} + \beta^{\mathrm{PA,IML}}_{ijkl}$ where $\beta^{\mathrm{PA,EL}}_{ijkl}, \beta^{\mathrm{PA,I2L}}_{ijkl}, \beta^{\mathrm{PA,IML}}_{ijkl}$ are the function $\beta^{\mathrm{PA}}_{ijkl}$ whose positions $i,j,k,l$ are in external loops, internal 2-loops, and internal multi-loops, respectively. The functions $\beta^{\mathrm{PA,EL}}_{ijkl}, \beta^{\mathrm{PA,I2L}}_{ijkl}, \beta^{\mathrm{PA,IML}}_{ijkl}$ can be computed based on inside partition functions $\{\alpha^{x}_{ijkl}, \alpha^{y}_{uv}\}$:

$$\begin{cases} \beta^{\mathrm{PA,EL}}_{ijkl} = \alpha^{\mathrm{EL,for}}_{i-1,k-1} \alpha^{\mathrm{EL,back}}_{j+1,l+1} \\ \beta^{\mathrm{PA,I2L}}_{ijkl} = \sum_{mnop} \beta^{\mathrm{PA}}_{mnop} \exp(\beta(\frac{p^{\mathrm{PA}}_{mnop}}{p^{\mathrm{PA,rand}}_{mnop}} - fe^{\mathrm{I2L}}_{mnij} - fe^{\mathrm{I2L}}_{opkl})) \alpha^{\mathrm{SA,for}}_{i-1,k-1} \alpha^{\mathrm{SA,back}}_{j+1,l+1} \\ \beta^{\mathrm{PA,IML}}_{ijkl} = \exp(-2\beta(fe^{\mathrm{IML,CBP}} + fe^{\mathrm{IML,ABP}})) \sum_{mnop} \beta^{\mathrm{PA}}_{mnop} \exp(\beta \frac{p^{\mathrm{PA}}_{mnop}}{p^{\mathrm{PA,rand}}_{mnop}})(\alpha^{\mathrm{left}}_{i-1,k-1} \alpha^{\mathrm{right}}_{j+1,l+1} + \alpha^{\mathrm{left\prime}}_{i-1,k-1} \alpha^{\mathrm{SA,back}}_{j+1,l+1}) \end{cases}$$

where $m < i < j < n, o < k < l < p$ and

$$\begin{cases} \alpha^{\mathrm{left}}_{i-1,k-1} = \alpha^{\mathrm{IML,for}}_{i-1,k-1} + \alpha^{\mathrm{1P,for}}_{i-1,k-1} + \alpha^{\mathrm{SA,for}}_{i-1,k-1} \\ \alpha^{\mathrm{right}}_{j+1,l+1} = \alpha^{\mathrm{IML,back}}_{j+1,l+1} + \alpha^{\mathrm{1P,back}}_{j+1,l+1} \\ \alpha^{\mathrm{left\prime}}_{i-1,k-1} = \alpha^{\mathrm{IML,for}}_{i-1,k-1} + \alpha^{\mathrm{1P,for}}_{i-1,k-1} \end{cases}.$$

(Supplementary figure 3)

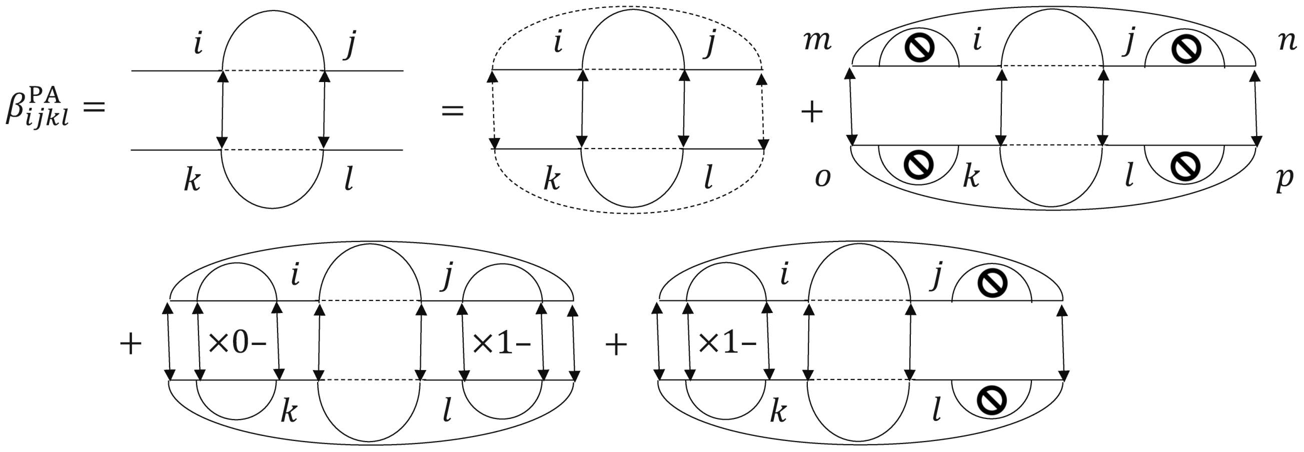


**Fig. 3.** An interpretation of the formulation of an outside partition function $\beta^{\mathrm{PA}}_{ijkl}$.
